# Molecular Translators as a Computational Primitive for Biomarker Discovery: Learnability Gains Under Conserved Information Ceilings

**DOI:** 10.64898/2026.04.27.720188

**Authors:** Payam Saisan, Sandip Pravin Patel

**Affiliations:** University of California, San Diego

## Abstract

Virtual molecular mapping systems such as MISO and GigaTIME introduce a potentially transformative primitive in computational pathology: translation of H&E whole-slide images into biologically structured molecular representations, learned on paired cohorts and deployed as an inference-time map. Despite sustained progress in machine learning, H&E-to-molecular-biomarker (e.g., gene mutation) prediction continues to exhibit recurrent field-level performance plateaus whose drivers remain poorly resolved. It remains unclear whether continued optimization targets a removable methodological limitation or instead presses against an intrinsic ceiling imposed by morphology. We develop a formal framework characterizing what deterministic translators can and cannot change. Histology-based biomarker modeling is governed by two constraints: method-limited gaps (finite labels, weak supervision, structured nuisance) and modality-limited ceilings (intrinsic slide-specific information in morphology). Because deterministic translation introduces no new slide-level measurements at inference, H&E information ceilings are conserved; however, translation can still improve finite-sample learnability, yielding an apparent information–performance paradox that we formalize as learnability gains under conserved information ceilings. We derive falsifiable signatures distinguishing these regimes and characterize them in controlled analytical experiments anchored to representative systems, including MISO and GigaTIME. We introduce an open-source toolkit comprising learning regime diagnosis, information-ceiling estimations, phase analyses, fidelity perturbation tests, and shortcut-confounding stress tests as an operational rubric for identifying and overcoming removable performance plateaus in translator-assisted molecular biomarker discovery and computational pathology.

## 1 Introduction: information ceilings, performance gaps and the information–performance paradox

Weakly supervised learning on hematoxylin and eosin (H&E) whole-slide images (WSIs) achieves strong performance on morphology-rich tasks such as tumor detection and grading (1). The field’s scope expanded at an early inflection point with (2), which showed that routine histology contains detectable molecular signal, including oncogenic alterations, without additional measurements at inference. Notably, that same result already exhibited the now-familiar gap: morphology-level discrimination reached histologic subtype AUC of approximately 0.97, whereas mutation prediction for common LUAD genes was substantially lower (AUCs of approximately 0.73–0.86). Since then, molecular alteration prediction from H&E has become a high-velocity empirical race involving larger encoders, larger cohorts, and stronger multiple instance learning (MIL) pipelines, where slide-level labels are observed but patch-level relevance remains latent (8; 9; 10). Yet for many molecular endpoints, performance repeatedly re-converges to a similar envelope across cohorts, institutions, and model families (2; 12; 13; 14; 15). This repeated pattern suggests a potentially systematic constraint rather than an idiosyncratic modeling failure.

EGFR mutation prediction in lung adenocarcinoma is a canonical example of this broader pattern. The literature supports a consistent picture: routine H&E contains nontrivial correlates of EGFR status (2), and while modern attention-based MIL, and foundation-model pipelines can improve prediction (13; 24; 23), performance often degrades under external validation and cohort heterogeneity. This is consistent with structured nuisance and dataset shift arising from site, stain, scanner, case mix, and histologic variability (14). Systematic review evidence further suggests that pooled performance remains substantially below that achieved on morphology-dominant diagnostic tasks (15). This motivates a basic question: are such empirical plateaus primarily *modality-limited*, reflecting an intrinsic H&E information ceiling, or *method-limited*, reflecting finite-label regimes, weak supervision, and structured nuisance that impede learning from the deployable H&E signal?

Many molecular alterations drive downstream transcriptional and proteomic programs that assays such as RNA sequencing, spatial transcriptomics, proteomics, and immunohistochemistry measure more directly, whereas H&E morphology provides an indirect projection of those states. Distinct molecular programs can therefore map to similar histologic appearances, as documented across driver-defined LUAD subtypes (31). Morphological representations are further perturbed by non-biological variation, including scanner, and tissue-processing effects (32), which can further decouple morphology from the underlying molecular state. By contrast, molecular assays measure products more tightly coupled to upstream biological programs. In LUAD, transcriptomic and proteomic measurements capture richer molecular-state structure than histologic appearance alone (29). EGFR-linked protein-expression programs exemplify this specificity (30). These are a few of many examples that motivate the central biological premise, not as a universal law, but as the specific regime considered in this paper; molecular measurements are more directly informative of molecular end points than H&E morphology:

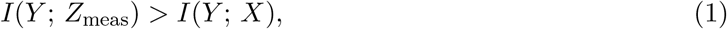

where *I*(·; ·) denotes mutual information, *X* is H&E morphology, *Y* is the molecular endpoint of interest, and *Z*_meas_ is a directly measured molecular map.

Although direct spatial molecular assays could be transformative for biomarker prediction, they remain too costly and throughput-limited for routine large-scale deployment relative to H&E. This deployment asymmetry creates the appeal of virtual molecular mapping: one may learn a translator *h*(·) on external paired cohorts and then deploy the frozen map 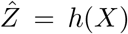 as an intermediate molecular representation derived from H&E alone. Recent systems such as MISO and GigaTIME, along with multi-omic and transcriptomics-mediated pipelines such as MOSBY (5) and ENLIGHT– DeepPT (6), make this possibility tangible, shifting the practical inference target from direct *X* → *Y* prediction toward deployed translation followed by downstream learning on 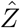.

A natural intuition here is to treat a high-fidelity translator as a virtual molecular measurement and expect the molecular advantage of Eq. (1) to carry over at deployment. But because 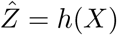 is a deterministic post-processing of *X*, it cannot introduce new slide-specific information about *Y*. This yields a seeming information–performance paradox: deterministic translation cannot increase information about *Y*, yet it can improve finite-sample discrimination:

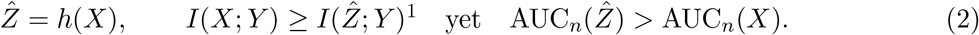

We resolve the tension expressed in Eq. (2) by separating two constraints on weakly supervised H&E biomarker modeling. First, an H&E deployment ceiling is set by the intrinsic information content of morphology and is conserved under deterministic translation. Second, a finite-sample method gap arises primarily from weak supervision and limited labeled sample size *n*. There is no contradiction: the practical advantage of translation is not additional slide-specific information at deployment, but population-level morphological structure acquired from external paired cohorts and encoded into *h*. In this sense, 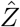 is best understood as a paired-data-learned prior that reshapes geometry and suppresses structured nuisance.

Critical to the analytical approach taken here is that this distinction is not identifiable from real H&E data alone. Because any empirical gap between *X* and 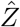 conflates the unknown Bayes-optimal ceiling from morphology with finite-sample learnability, real cohorts cannot disentangle the two. Nor do they allow nuisance structure and translator fidelity to be varied independently while holding the underlying task fixed. We therefore establish our core claims in a controlled synthetic setting where these quantities are explicit by construction, allowing convergence behavior to be interpreted against known ground truth.

The relevant object of study here is broader than a single translator, but a space of paired-data-learned maps from morphology into biologically structured intermediate representations. Even when the information ceiling is preserved, these translators need not be equivalent: some may be neutral or harmful, while others substantially reduce the labeled cohort size needed to learn a biomarker from routine histology. Since deterministic translation cannot introduce new information, the practical problem is to identify representations that reorganize existing morphological information into a form that improves finite-*n* learnability^2^ in weakly supervised, heterogeneous settings. Figure 2 summarizes the baseline weakly supervised pipeline and the expected finite-*n* benefit of translator-assisted prediction under a conserved H&E ceiling.

**Figure 1:**
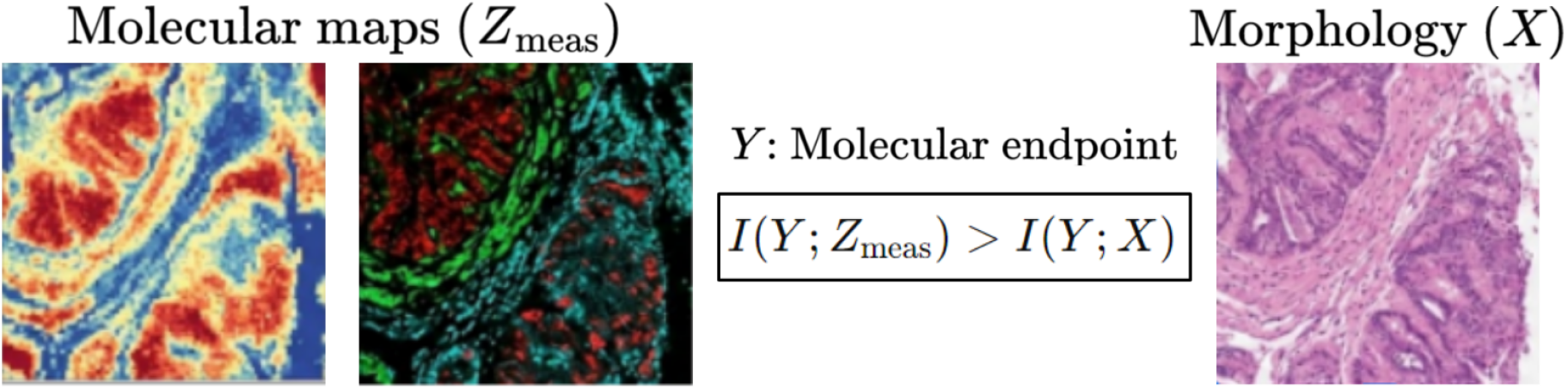
Assumed biological regime: Directly measured molecular maps *Z*_meas_ (e.g., spatial RNA or protein) carry more information about the molecular endpoint *Y* than deployable morphology *X*. This motivates *paired-data-trained cross-modal translators* 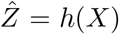 as proxies for a biologically privileged modality.

**Figure 2:**
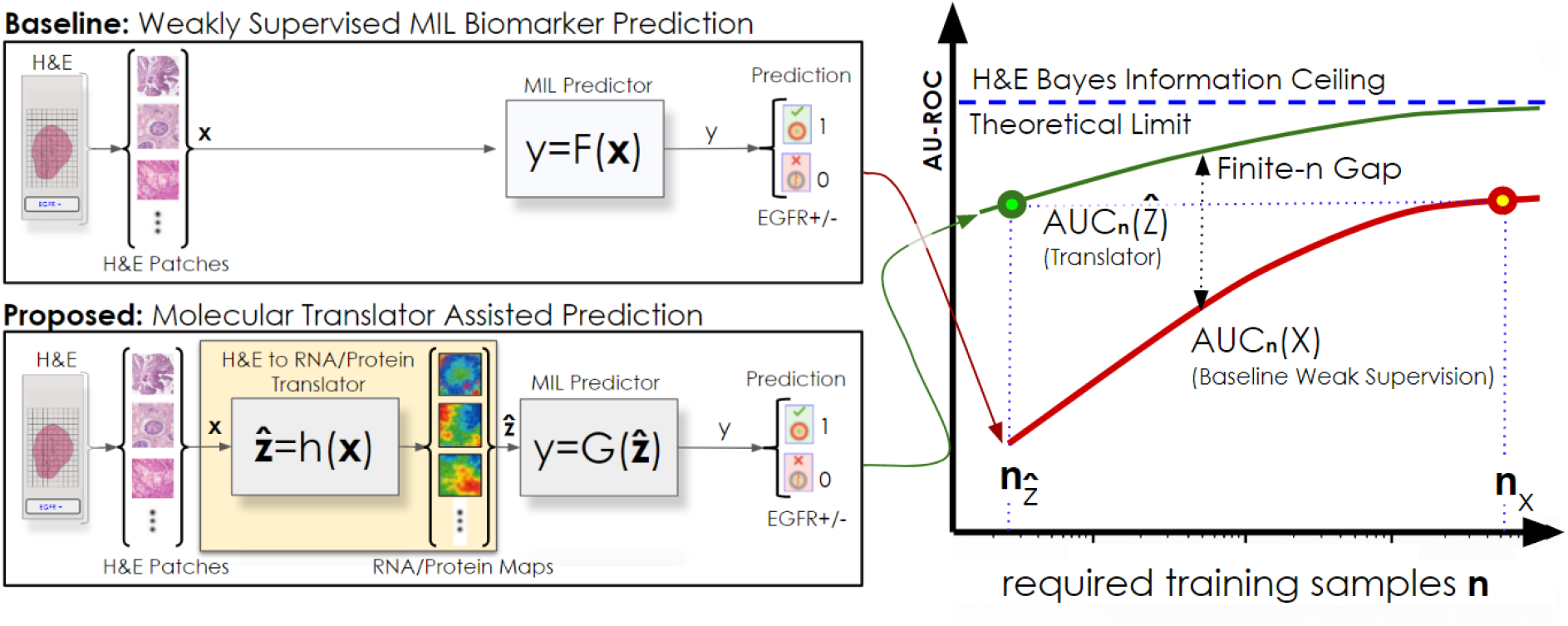
Translator-assisted prediction and finite-*n* gains under conserved ceilings. **Left:** Baseline weakly supervised biomarker prediction from H&E patches, **y** = *F* (**x**), versus translator-assisted prediction 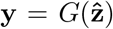 with 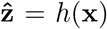, where the translator is learned on paired cohorts and frozen at deployment (depicting individual-sample instance flow for **x**). **Right:** At the population level, deterministic translation preserves the H&E deployment ceiling, but can reduce the finite-sample method gap, yielding 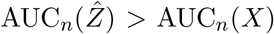; equivalently, fewer labeled training samples are required to reach a comparable performance level 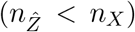 in practical weak-supervision regimes.

Beyond the central premise that transformed representations can improve prediction in low-data regimes, this work brings into focus 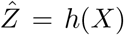 as a specific deployment primitive for biomarker learning: it imports population-level morphological structure learned from paired cohorts into an H&E-only setting without adding new slide-specific information at inference. Our goal is to provide a diagnostic framework for determining when translation reduces a removable method gap and when a recurrent plateau is modality-limited. We provide (i) a ceiling-containment result showing 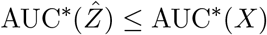 for any deterministic deployed translator, (ii) a finite-*n* learnability formalization that explains how 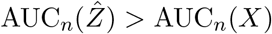 can occur despite deployment-time ceiling non-expansion, and (iii) falsifiable signatures, low-*n* concentration, heterogeneity dependence, and fidelity dependence, that provide a diagnostic rubric for whether observed gains are consistent with a learnability-mediated method gap vs. a modality-limited ceiling. Translation, however, also leverages an additional resource: external paired cohorts of size *n*_pair_ used to learn *h*. Gains in 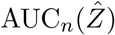 should therefore be interpreted as gains from importing a paired-data-learned prior, and compared against resource-matched alternatives when making engineering claims.

### 1.1 Deployment modes and scope

Our theory is translator-agnostic: it applies to any deterministic deployed representation 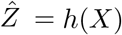, regardless of architecture or paired assay. We distinguish two deployment modes. In *Mode A* (the focus here), a translator computes 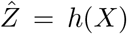 from H&E at inference, and downstream prediction is performed on 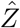(optionally fused with *X*). In *Mode B*, molecular information is used only during training to shape an H&E-only predictor that is deployed without computing 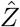 at inference. This setting is closer to learning using privileged information or generalized distillation than to deployment-time translated inference; recent pathology work such as TriDeNT follows the same training-only pattern (7; 11; 36). We do not consider Mode B further here. Table 2 situates the present study within the broader class of deployed H&E→molecular translation systems that instantiate Mode A. In Section 2.3, we focus on GigaTIME (protein) and MISO (spatial transcriptomics) as two of the most successful exemplars in their respective modalities, which serve as empirical anchors for our analytical framework.

**Table 1:** Landmark results in weakly supervised whole-slide image learning. Representative studies illustrating the contrast between morphology-driven diagnostic tasks and molecular or microenvironmental biomarker inference from routine H&E histology.

| Study | Year | Task | Target | AUC performance |
| --- | --- | --- | --- | --- |
| Campanella et al. (1) | 2019 | Diagnosis / detection | Cancer, metastasis | > 0.97 |
| Coudray et al. (2) | 2018 | Mutation prediction | EGFR, KRAS, others | ~0.73–0.86 |
| Kather et al. (12) | 2019 | Molecular phenotype | MSI | ~0.77–0.84 |
| Pao et al. (14) | 2023 | Mutation prediction | EGFR (LUAD) | ~0.85, one-center |
| Nguyen et al. (15) | 2025 | Review | EGFR (meta-analysis) | ~0.75, pooled |

Reported AUCs are not strictly comparable across studies due to cohort composition, endpoints, preprocessing, and evaluation protocols; values are included to illustrate the persistent diagnostic-vs-molecular performance gap.

**Table 2:**
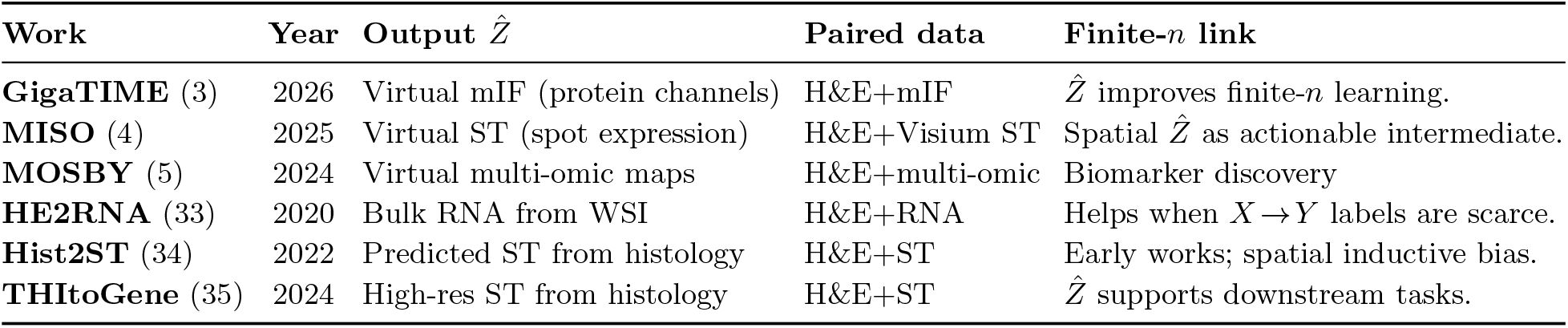
Representative H&E → molecular translation systems producing intermediate representations 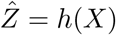.

## 2 Information limits and finite-sample learnability

### 2.1 Deterministic maps cannot introduce new information

Let *X* denote deployable H&E-derived input available at inference (e.g., slide or patch representations) and *Y* a target label (e.g., EGFR mutation status). Deterministic deployed translation computes

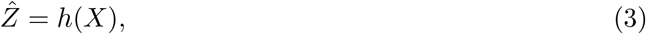

where *h*(·) is learned from external paired data (e.g., H&E with measured molecular maps) and then frozen at deployment.

We distinguish two limiting factors in weakly supervised biomarker modeling from H&E. *Method-limited* settings arise when finite labeled cohort size *n* induces large estimation error due to high-dimensional inputs, structured nuisance variation (site, stain, scanner, processing, case mix), and weak localization. *Modality-limited* settings arise when, even with unlimited labeled data and optimal inference, H&E does not uniquely identify the target because distinct molecular states can yield similar morphology.

Define the Bayes-optimal deployment ceiling for a representation *R* as

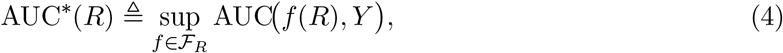

where *ℱ*_*R*_ denotes all measurable scoring rules based on *R*. For a fixed learning pipeline trained with *n* labeled samples, let AUC_*n*_(*R*) denote its expected test AUC. By construction,

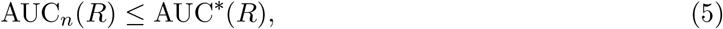

with equality only in idealized limits. Method-limited regimes correspond to a large gap AUC^*^(*R*) AUC_*n*_− (*R*); modality-limited regimes correspond to AUC^*^(*R*) itself being bounded away from 1.

Because 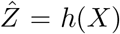 is deterministic, every predictor based on 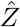 can be written as a predictor based on *X*:

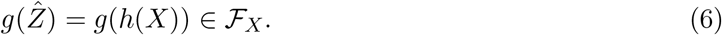

Thus the predictor class induced by 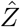 is a subset of the predictor class induced by *X*, implying

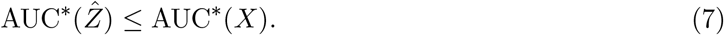

Translation preserves or lowers the deployment ceiling but cannot raise it. Equality can occur if *h*(*X*) is sufficient for *Y* under the deployment distribution, but it cannot be exceeded.

#### The finite-sample paradox

Ceiling ordering is asymptotic. In realistic method-limited regimes, observed performance is dominated by estimation error and inductive bias rather than by the ultimate deployment ceiling. It is therefore entirely possible that

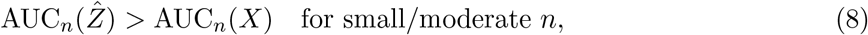

while still respecting Eq. (7). Fig. 3 demonstrates this in a linear benchmark: prediction from 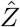 substantially outperforms direct prediction from *X* at low label budget, with the gap shrinking as *n* increases. Performance from *X* also degrades under nuisance whereas 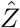 remains comparatively stable, showing that learnability gains concentrate at low *n* and amplify under structured heterogeneity.

**Figure 3:**
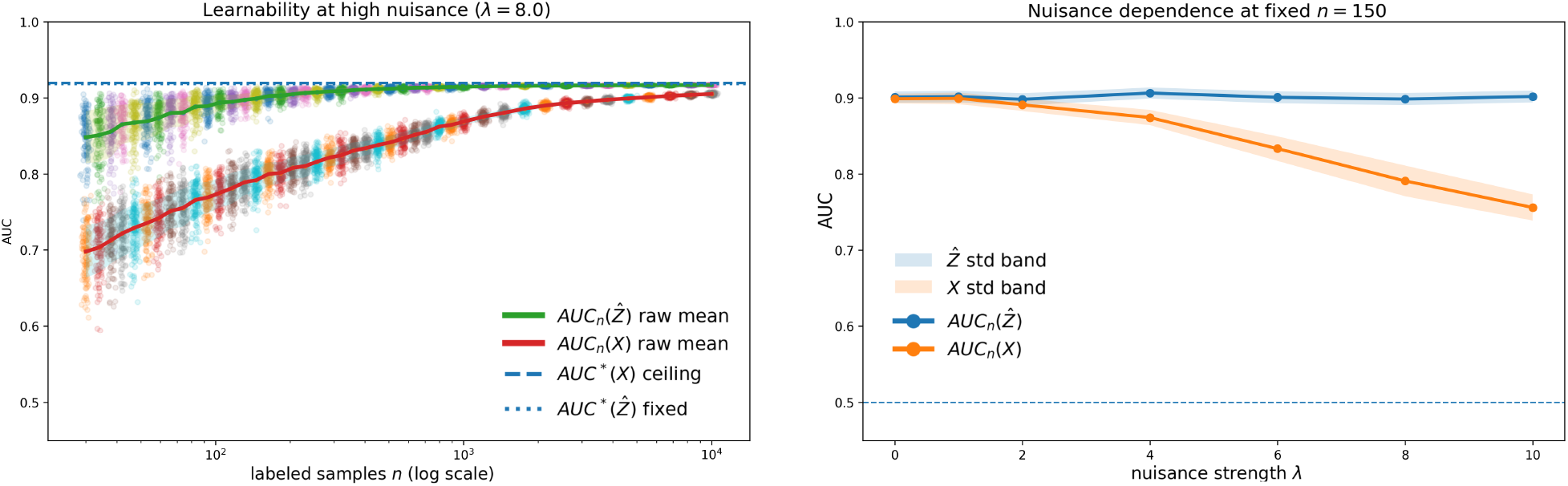
Linear signatures of finite-sample learnability gain under structured nuisance. **Left:** low-*n* concentration. At fixed high nuisance, prediction from 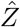 substantially outperforms direct prediction from *X* at small labeled sample sizes *n*_label_, with the gap shrinking as *n*_label_ increases. **Right:** nuisance dependence at fixed *n*_label_. Performance from *X* degrades as nuisance strength *λ* increases, whereas learning from 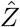 remains comparatively stable. In this linear benchmark, the two approaches approach similar large-*n* performance, consistent with conserved deployment ceilings.

### 2.2 Geometry shaping and structured nuisance suppression

We model translation as an externally learned prior that reshapes representation geometry. In the intended pipeline:

1. Learn *h*(·) from paired data 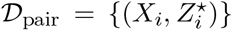, where *Z*^⋆^ denotes the directly measured paired biological modality, and freeze it.
2. Train a downstream predictor of *Y* using labeled data *𝒟*_label_ = {(*X*_*j*_, *Y*_*j*_)}, with inputs either 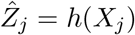

Because the translator is learned from paired data, it can encode population-level morphology– biology structure that is hard to infer from *Y* labels alone. If *h*(·) suppresses nuisance directions or reduces effective dimension while retaining label-relevant structure, it can reduce sample complexity and improve performance at fixed *n* in method-limited regimes (Eq. (8)).

#### Falsifiable signatures

This hypothesis yields three testable predictions:

- **Low-***n* **concentration:** gains 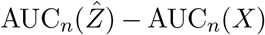 are largest at small or moderate *n* and shrink as *n* increases. In favorable cases the two approaches converge toward similar large-*n* performance; under lossy translation, 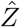 may instead retain a slightly lower large-*n* ceiling (Figs. 3 and 7).
- **Heterogeneity dependence:** gains increase with structured nuisance or heterogeneity that inflates estimation error in *X*-space (Figs. 6 and 8).
- **Fidelity dependence:** gains degrade monotonically when the translated representation is explicitly degraded, for example by additive corruption of 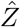(Fig. 6).

These signatures induce a **phase structure**: gains peak in the low-label, high-nuisance, high-fidelity regime and weaken outside it, as demonstrated in controlled proof-of-concept experiments that sweep label budget, nuisance strength, and fidelity (Fig. 6). The phase structure clarifies when translation is worth optimizing. One should optimize the translator when learning curves remain steep in the low-label regime, gains amplify under nuisance or heterogeneity, and improvements in fidelity improve downstream performance. One should stop when gains collapse toward zero despite increasing labels or improving fidelity, or when direct prediction from *X* catches up at large *n*, suggesting proximity to a modality-limited ceiling. Table 3 extends these signatures into a diagnostic workflow for deciding whether to invest in labels, improve *h*(*X*), strengthen paired supervision, or stop H&E-only iteration.

**Table 3:** Workflow for plateau diagnosis and next steps. Signatures for deciding whether to invest in labels, improve *h*(*X*), strengthen paired supervision, or stop H&E-only iteration.

| Signature | Observation | Diagnosis | Next Step |
| --- | --- | --- | --- |
| **Label sensitivity** | $AUC_n$ improves significantly with increasing $n_{\text{label}}$ (not flat); gains are largest at low $n_{\text{label}}$. | Method-limited (*labels / sample complexity*) | Expand or rebalance the $Y$-labeled cohort; if limited labels, prioritize learning $\hat{Z}$ to reduce label demand. |
| **Nuisance stress / shift** | Out-of-site or site-stratified evaluation shows strong degradation for $X$; $\hat{Z}$ is more stable and the gap grows with heterogeneity ($\lambda$). | Method-limited (*structured nuisance / shift*) | Confirm the pattern under deployment-like splits; then improve $h(X)$ for invariance and normalization before adding end-to-end model complexity. |
| **Fidelity sensitivity** | Improving paired supervision (larger $n_{\text{pair}}$, better assays) increases $AUC_n(\hat{Z})$ monotonically under a fixed downstream protocol. | Method-limited (*representation quality*) | Strengthen paired supervision and calibration; then rerun the same downstream protocol to verify that gains rise with translator fidelity. |
| **Ceiling convergence** | Performance is insensitive to both $n_{\text{label}}$ and translator fidelity; $\Delta AUC = AUC_n(\hat{Z}) - AUC_n(X) \approx 0$ and both curves flatten. | Consistent with modality-limited ceiling (*after sanity checks*) | Rule out leakage, site-split artifacts, and shortcut effects; if curves remain flat, stop H&E-only iteration and use different endpoint/modality. |
| **Shortcut / circularity check** | Gains appear under random splits but vanish or reverse under strict site-stratified splits and/or fail to decay under controlled fidelity degradation. | Shortcut / circularity risk (*confounding or evaluation artifact*) | Lock model selection and enforce deployment-like evaluation; then run fidelity-degradation and calibration checks before treating any gain as biology-aligned. |

**Notation:** $n_{\text{label}}$ = labeled training size; $n_{\text{pair}}$ = paired cohort size for learning $h$ ; $\lambda$ = nuisance / heterogeneity strength; $AUC_n(\cdot)$ = expected test AUROC under a fixed downstream protocol; $\Delta AUC = AUC_n(\hat{Z}) - AUC_n(X)$ .

**Table 4:** Mapping GigaTIME objects into our framework (Mode A).

| Framework object | GigaTIME instantiation |
| --- | --- |
| $X$ (deployable morphology) | H&E whole-slide imagery (or patches/features), available at inference. |
| $Z^*$ (paired measured target for learning $h$) | Measured multiplex immunofluorescence (mIF) channels on paired tissue. |
| $h(\cdot)$ (translator) | Supervised H&E→mIF translator trained on paired H&E+mIF data, then frozen. |
| $\hat{Z} = h(X)$ (deployed representation) | Virtual mIF channels generated from H&E at inference (multi-channel intermediate representation). |
| Downstream $Y$ | Biomarker association / stratification / survival endpoints (population-scale analyses). |

**Table 5:** Mapping MISO objects into our framework (Mode A) with an explicit ceiling proxy.

| Framework object | MISO instantiation |
| --- | --- |
| $X$ (deployable morphology) | H&E whole-slide imagery (patches/features), available at inference. |
| $Z^*$ (paired measured target for learning $h$) | Spatial transcriptomics counts / gene expression maps on paired tissue. |
| $h(\cdot)$ (translator) | Supervised H&E→spTx predictor trained on paired data, then frozen. |
| $\hat{Z} = h(X)$ (deployed representation) | Predicted spatial gene expression maps (or derived representations) computed from H&E at inference. |
| Ceiling proxy | $R_{\max}$: gene-wise maximum achievable correlation with noisy observed counts under a count-noise model. |

### 2.3 Empirical anchors: GigaTIME and MISO

We use two deployed Mode A exemplars as empirical anchors rather than as controlled empirical tests of the present framework. GigaTIME is most informative about heterogeneity and cross-cohort shift under deployment-like nuisance, whereas MISO is most informative about ceiling-limited behavior and finite-sample learnability under a fixed deployment modality.

#### GigaTIME as deployed translation 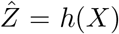

GigaTIME (3) learns a deterministic translator from routine histology *X* (H&E) to a multi-channel virtual multiplex immunofluorescence representation 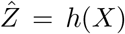 and deploys 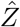 at inference for downstream association, stratification, and survival analyses.

GigaTIME does not report AUROC for a fixed clinical label *Y* under a controlled *X* versus 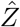 comparison. Instead, it provides three quantitative anchors that map to the present framework:

1. **Heterogeneity stress-test**. Deployed on a Providence cohort (14,256 H&E WSIs; 51 hospitals; *>* 1,000 clinics; 24 cancer types / 306 subtypes) and compared with an external TCGA virtual population (10,200 tumors), aggregated virtual protein activations show cross-cohort concordance (Spearman = 0.88 across cancer subtypes) and overlapping biomarker associations (Fisher *p <* 2 *×* 10^−9^; 80 shared).
2. **Translator fidelity**. On paired H&E+mIF evaluation, GigaTIME outperforms CycleGAN on 15/21 channels; naive baselines fail (for example, DAPI Dice 0.12 versus 0.72), establishing that 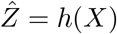 is a measurable, non-degenerate transformation.
3. **Downstream discrimination**. Virtual protein activations support survival stratification; a composite 21-channel signature stratifies more strongly than individual channels, consistent with finite-sample learnability under conserved ceilings.

Within the present framework, GigaTIME anchors the *heterogeneity-dependence* signature.

#### MISO and the gene-wise ceiling *R*_max_

MISO (4) predicts spatial transcriptomics from histology as 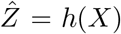 with no spatial assay at inference, satisfying Mode A deployment. It is well aligned with the present framework because it introduces an explicit gene-wise ceiling proxy *R*_max_, providing a measurable axis for modality-limited behavior while remaining H&E-only at deployment.

MISO models observed spot-wise counts for gene *g* as noise-corrupted observations of latent rates, e.g.,

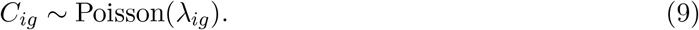

Even an oracle that predicts the true latent rates *λ*_*ig*_ cannot correlate perfectly with the noisy observed counts *C*_*ig*_. MISO defines the resulting upper bound on achievable Pearson correlation for each gene as

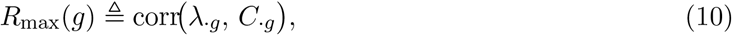

and uses *R*_max_ as an explicit ceiling proxy to interpret which genes are intrinsically modality-limited. MISO provides three quantitative anchors that map cleanly onto the present framework:

1. **Explicit ceiling proxy**. *R*_max_ supplies a gene-wise ladder of achievable performance under measurement noise; empirical results track this axis, supporting the idea that some targets are intrinsically ceiling-limited under histology-only deployment.
2. **Learnability gains**. Knowledge distillation improves spot-level prediction (Pearson 0.369 → 0.427 and Spearman 0.371 → 0.430, with bootstrapped 95% confidence intervals), consistent with improved learnability under fixed deployment modality.
3. **Downstream discrimination**. The survival-risk model achieves a cross-validated concordance index of 0.67, showing ranking-style downstream prediction from histology-derived representations even when deployment remains H&E-only.

Within the present framework, MISO anchors the *ceiling-axis* signature via *R*_max_ and provides concrete evidence that representation and learning choices can move finite-sample performance while the deployment modality remains fixed.

GigaTIME and MISO show that the abstract regime analyzed here already exists in deployed form. However, neither study systematically varies label budget (*n*_label_), structured nuisance complexity (*λ*), and translation fidelity (*n*_pair_, corruption) in the controlled manner required to test the signatures above. Our proof-of-concept analytic experiments (Section 3) fill this gap by isolating mechanism rather than merely establishing the existence of the effect.

## 3 Controlled analytical characterization of finite-Sample learnability gains

The previous section identified three signatures of learnability-mediated gain under conserved deployment ceilings: low-*n* concentration, heterogeneity dependence, and fidelity dependence. Establishing these signatures requires independent control of the morphology-limited ceiling, structured nuisance, and translator fidelity. Real H&E cohorts cannot provide such control. In real studies, the observed gap between learning from *X* and from 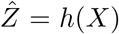 entangles unknown ceiling effects with finite-sample learnability, while paired-data quality, nuisance structure, and task difficulty co-vary in ways that cannot be cleanly separated. The synthetic framework is therefore used here because it is the only setting in which all components of the mechanism are simultaneously identifiable: the morphology-limited ceiling, unobservable in any real cohort, is known by construction, each factor can be perturbed independently, and convergence can be interpreted against explicit ceilings. The aim is not to mimic any particular assay or disease setting, but to isolate the regime in which deterministic translation improves learnability while leaving deployment-time information unchanged.

Throughout this section, *B* denotes latent biology, *X* the deployable representation available at inference, and *Z*^⋆^ the paired biological target used to train the translator. The translator is learned from paired data, frozen, and then compared downstream against direct learning from *X*. Standardized *ℓ*_2_-regularized logistic regression is used as a deliberately simple fixed probe, so differences should be interpreted as differences in finite-sample learnability under a common protocol rather than as the result of downstream architecture search (37).

At a schematic level, the deployable representation mixes biological signal with structured nuisance,

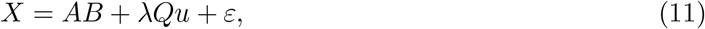

where *u* denotes nuisance variation, *A* mixes latent biology into the observed space, *Q* spans a nuisance subspace, and *λ* controls nuisance strength. The question is whether a translator learned from paired data can reorganize this same deployment-time information into a representation that is easier to learn from when labels are scarce.

We answer that question in four steps. We first verify the two sanity checks implied by deterministic translation: mutual information and Bayes-optimal deployment ceilings for 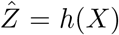 cannot exceed those of *X*. We then analyze an exactly tractable linear-Gaussian benchmark, move to a nonlinear latent world in which translation can be simultaneously useful and lossy, and finally summarize robustness sweeps over translator fidelity, nuisance entanglement, and paired-data availability.

### 3.1 Sanity checks for deterministic translation

In the linear-Gaussian benchmark, a ridge translator is relearned at each paired-cohort size *n*_pair_ and then frozen. Figure 4 confirms the two constraints established in Section 2: as translator quality improves, 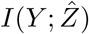 increases but remains below *I*(*Y* ; *X*), and 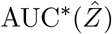 approaches but does not exceed AUC^*^(*X*). Any downstream advantage of 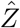 must therefore arise through finite-sample learnability rather than new deployment-time information.

**Figure 4:**
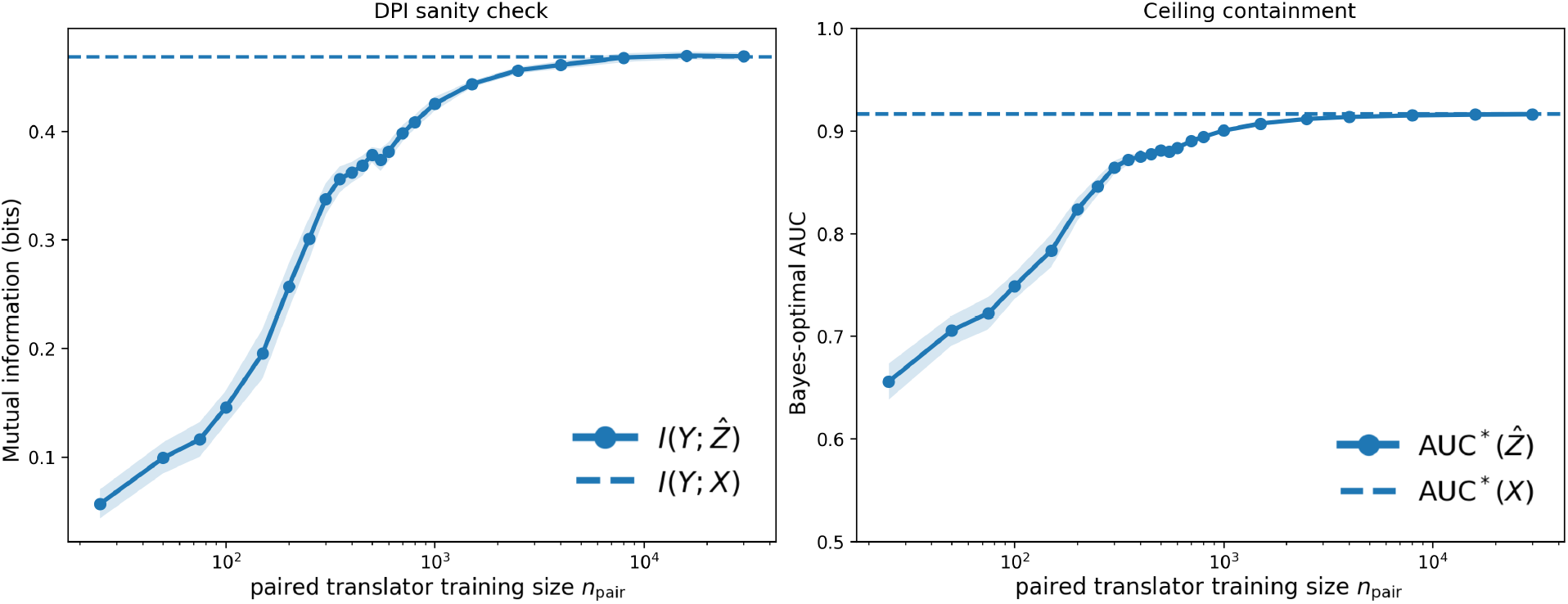
Sanity checks for deterministic translation in the linear-Gaussian benchmark. For a fixed deployment distribution and deployable representation *X*, a ridge translator 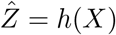 is relearned at each paired-cohort size *n*_pair_. Horizontal reference lines show the fixed deployable information and Bayes ceiling for *X*; curves show the corresponding quantities for 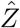. **Left:** 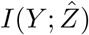 increases with translator quality but remains below *I*(*Y* ; *X*), consistent with the data-processing inequality. **Right:** 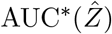 rises toward but does not exceed AUC^*^(*X*). Any downstream gain from 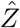 must therefore reflect finite-sample learnability rather than new deployment-time information.

**Figure 5:**
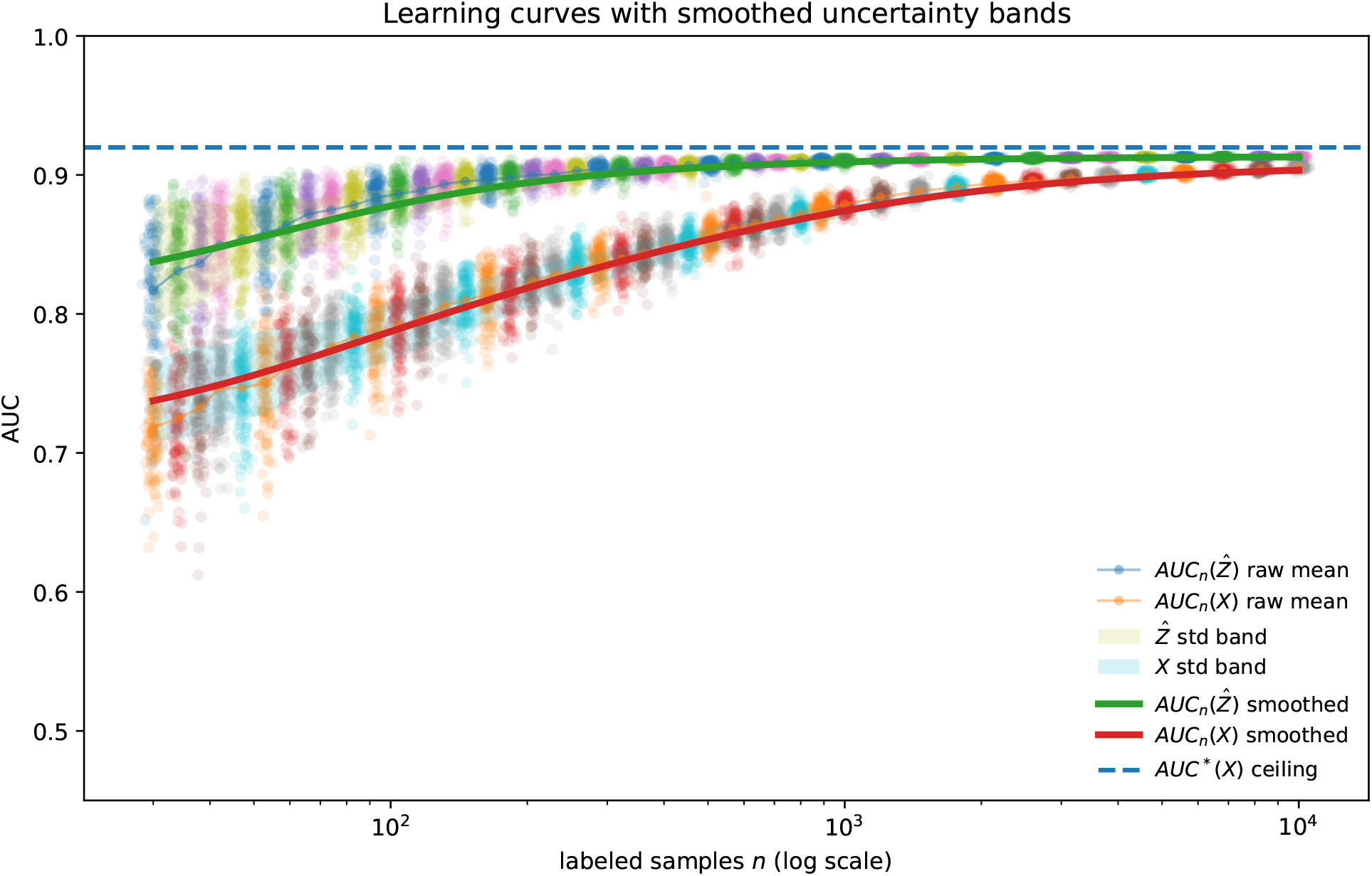
Linear benchmark: finite-sample learnability gains under structured nuisance. Learning curves for prediction from 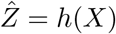 and from the raw deployable representation *X* in the linear-Gaussian benchmark. Points denote replicate means, shaded bands denote uncertainty, and the dashed line marks the Bayes-optimal ceiling in *X*-space. Although deterministic translation cannot increase the asymptotic information ceiling, the translated representation yields substantially better predictive performance in the low-label regime, consistent with improved finite-sample learnability under structured nuisance.

### 3.2 The linear regime: isolating learnability from information

We begin with a setting that is simple enough to analyze exactly yet rich enough to express the paper’s core mechanism: class signal lives in low-dimensional latent biology, the paired target is comparatively well aligned with that biology, and the deployable representation is contaminated by structured nuisance.

#### Generative chain

Let *s* ∈ {−1, +1} with equal probability and *Y* = (*s* + 1)*/*2. The latent biological state *B* ∈ ℝ^*m*^ is class-conditionally Gaussian,

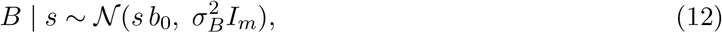

where *b*_0_ ∈ ℝ^*m*^ is a fixed biology-direction vector and 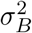 controls within-class biological variability, so the two classes differ only through a shift in the latent biological mean. The paired target used to train the translator is a noisy, biology-aligned readout

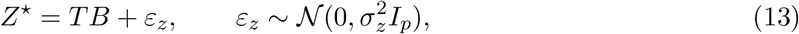

where *T* ∈ ℝ^*p×m*^ is chosen to be *identity-like*—the first min(*p, m*) biology axes are approximately preserved up to small random perturbation—modeling a paired assay that is well aligned with the biologically relevant factors without being an exact noise-free readout of *B*. The deployable representation available at inference is

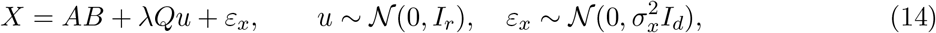

where *A* ∈ ℝ^*d×m*^ mixes biological signal into the observed space, *Q* ∈ ℝ^*d×r*^ has orthonormal columns spanning an *r*-dimensional nuisance subspace, and *λ* ≥ 0 controls nuisance strength. Combining (12) and (14) gives

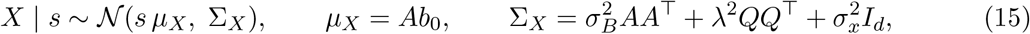

so class information appears entirely as the mean shift *±µ*_*X*_, while nuisance inflates covariance in the *Q*-subspace.

Given paired samples 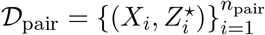, we fit a ridge translator

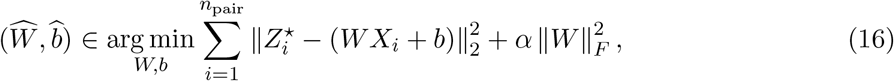

and freeze the affine map 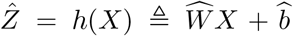. Paired data are used only to learn *h*; all downstream comparisons are between prediction from *X* and from 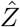 under the same deployment modality. Because 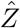 is an affine transform of a Gaussian, it is class-conditionally Gaussian with

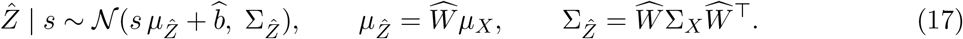

The intercept shifts both classes equally and does not affect AUROC.

#### Asymptotic ceilings and their containment

For balanced Gaussian classes with means *±µ* and shared covariance Σ, the Bayes-optimal score is linear, 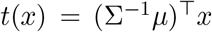, and the squared Mahalanobis separation determines the asymptotic ceiling via 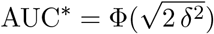. Defining

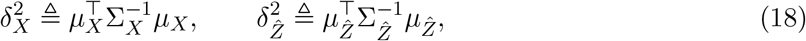

the two Bayes-optimal AUROCs are

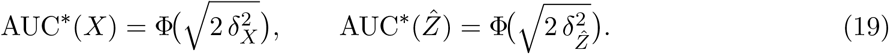

Because 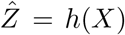 is deterministic after translator training, no deployment-time information is added, and the predictor class containment established in Section 2.1 directly implies

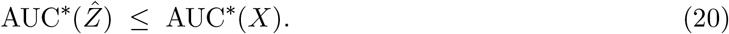

In this benchmark the paired target is biology-aligned, so 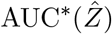 can approach AUC^*^(*X*) closely, but equality is not imposed by the construction. This is the structural point: any downstream advantage of 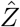 must arise not from additional information, but from more favorable finite-sample conditioning.

#### Finite-sample comparison

The point of the benchmark is not the ceiling but the low-label regime. Using only *n*_label_ labeled samples, we train one standardized *ℓ*_2_-regularized logistic regressor on (*X*_*j*_, *Y*_*j*_) and another on 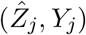, then evaluate both on a large held-out test set to obtain 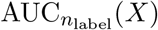 and 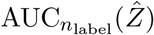. Regularization strength is set separately for the two representations rather than forced to be numerically identical, reflecting their different effective dimensions and conditioning. In the low-label experiment, nuisance is held fixed at a large value and *n*_label_ is varied with a single frozen translator. In the complementary nuisance experiment, *n*_label_ is held fixed while *λ* is varied, again with the translator frozen after training at a reference nuisance level. These experiments isolate the central mechanism: although deterministic translation cannot raise the asymptotic deployment ceiling, it can improve finite-sample learnability by presenting a more biology-aligned and less nuisance-dominated representation to the downstream learner.

These gains are not generic consequences of translation. When task-relevant structure is progressively removed from 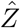, equivalently, when the translator is learned with weaker paired supervision, the low-label advantage decays monotonically toward the direct-*X* baseline. Across the label–nuisance plane, positive ΔAUC localizes to the low-label, high-nuisance corner and shrinks as label budget grows or nuisance weakens. Figure 6 makes this phase structure explicit and already points toward the broader lesson of the section: this is a finite-sample conditioning effect, not an asymptotic information gain.

**Figure 6:**
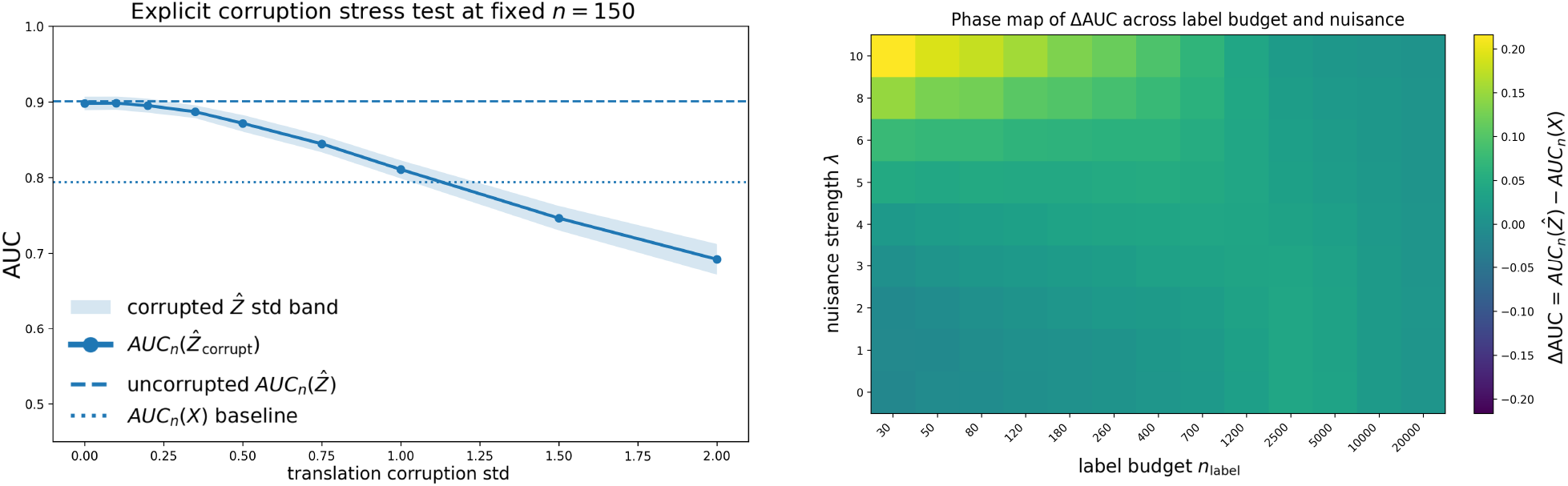
Fidelity dependence and phase structure of translator gains. **Left:** Downstream AUC as 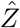 is progressively corrupted, reducing preservation of task-relevant biological structure. The advantage of 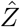 decays monotonically toward the direct-*X* baseline, confirming that the gain depends on translation fidelity. **Right:** heatmap of 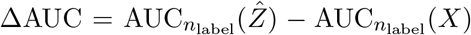 over label budget and nuisance strength. Gains localize to the low-label, high-nuisance regime and shrink as label budget increases or nuisance weakens, consistent with a finite-sample learnability effect under conserved deployment-time information.

### 3.3 Nonlinear extension: finite-sample gains under lossy translation

The linear-Gaussian benchmark isolates the mechanism in an analytically controlled setting, but real deployable representations are rarely linear or nuisance-free. We therefore move to a nonlinear latent world in which labels depend on a nonlinear function of latent biology, the paired modality remains comparatively aligned with the task-relevant axes, and the deployable representation is a higher-dimensional, nuisance-entangled observation of the same latent state.

#### Generative chain

Let *B* ∈ ℝ^*m*^ denote a latent biological state. Drawing two orthonormal directions *b*_0_, *b*_1_ ∈ ℝ^*m*^, we define

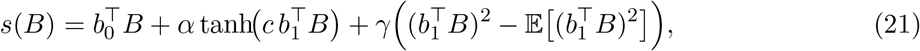

and set *Y* = 𝕀 [*s*(*B*) *>* median(*s*(*B*))] so that classes remain approximately balanced. An optional label-flip probability imposes a subunit ceiling representing irreducible mismatch between latent biology and the observed label.

The paired target *Z*^⋆^ ∈ ℝ^*p*^ is generated as a relatively direct nonlinear readout of *B*,

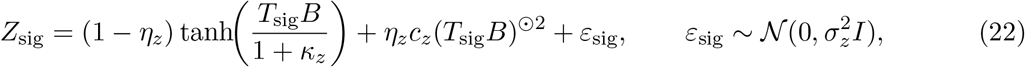

with optional weaker nuisance coordinates appended; their detailed form is not material to the argument. The key point is that *Z*^⋆^ remains comparatively well aligned with the latent axes that determine *Y*. The deployable representation is generated from the same latent state through a less favorable observation model,

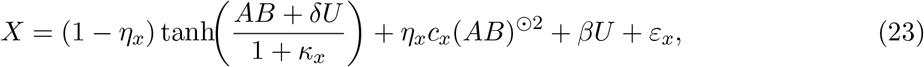

where *A* ∈ ℝ^*d×m*^ is a random mixing matrix, *U* ∈ ℝ^*d*^ is a structured low-rank nuisance term, and 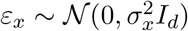. Relative to *Z*^⋆^, *X* is therefore higher-dimensional, more nuisance-entangled, and more strongly distorted by nonlinear compression and quadratic mixing.

#### Useful-yet-lossy regime

Given paired samples 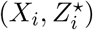, we fit a ridge translator *h* : ℝ^*d*^ → ℝ^*p*^, freeze it, and compare 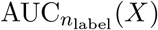 against 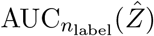 across labeled sample sizes. What the nonlinear construction adds is a regime in which early learnability gain and lower large-*n* performance coexist cleanly: 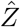 is helpful at low *n*_label_ because its geometry is easier to learn from, yet lossy enough that direct prediction from *X* can eventually catch up or slightly exceed it at large *n*_label_. Figure 7 shows exactly this regime.

**Figure 7:**
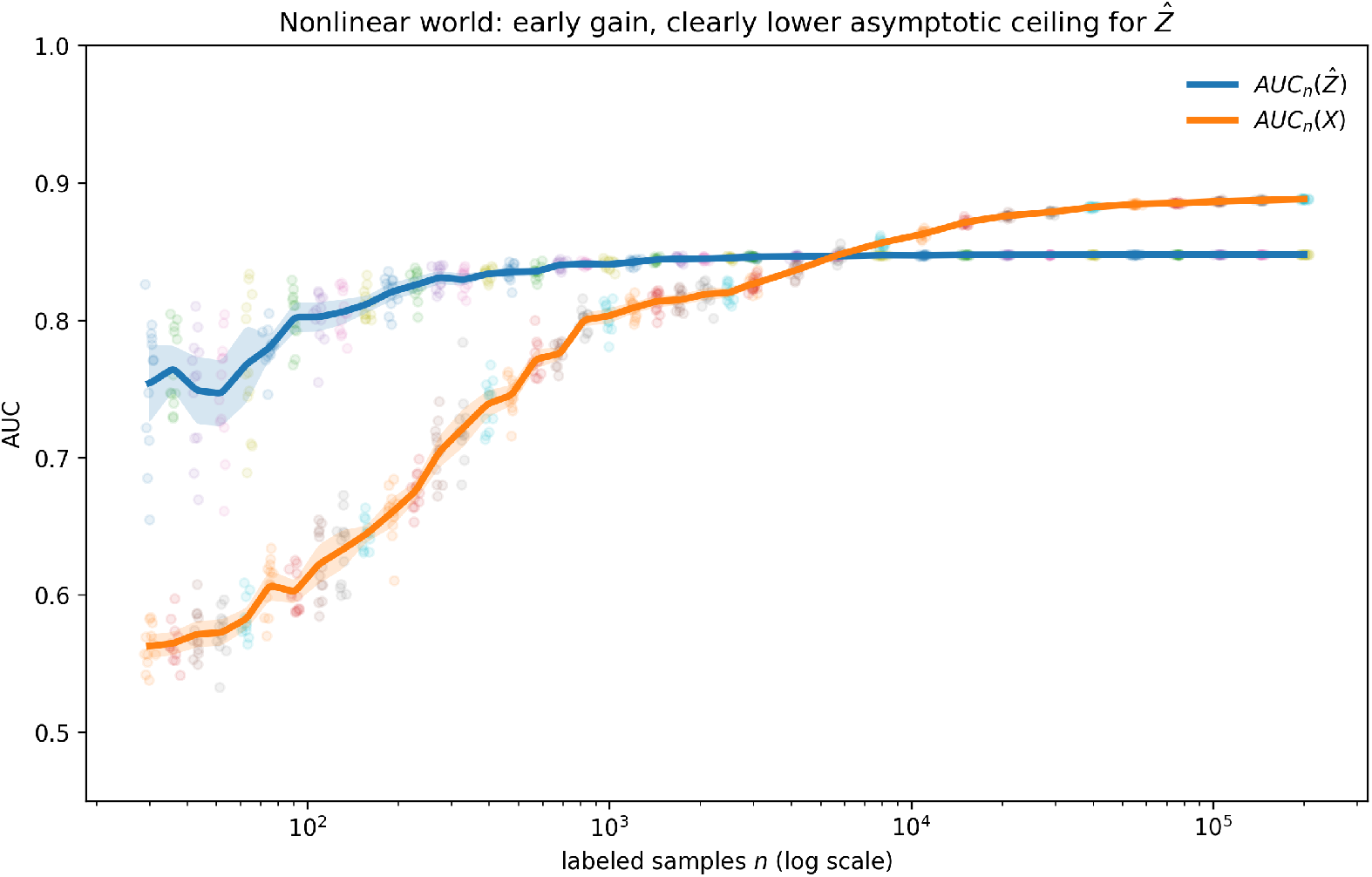
Nonlinear example: early learnability gain with lower large-*n* tail performance for 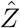. Learning curves for prediction from *X* and from 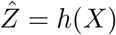 in a nonlinear latent-world simulation. The translated representation (blue) yields a strong advantage at small labeled sample sizes, while direct prediction from *X* (orange) catches up only at much larger sample sizes and attains slightly higher tail performance over the simulated label range, consistent with lossy translation.

### 3.4 Robustness of learnability gains across nonlinear sweeps

To determine whether this nonlinear effect is structural rather than tied to a single tuned example, we performed three sweep families: translator lossiness, nuisance entanglement in the deployable representation, and paired-sample availability for learning the translator. These correspond to three practical questions: how much task-relevant structure the learned map preserves, how difficult the raw deployable representation is to use directly, and how much paired data is needed to train a useful translator.

In the lossiness sweep, we varied paired-modality noise, translated dimensionality, translator regularization, and paired-data volume. In the nuisance sweep, we varied nuisance rank, amplitude, and nonlinear distortion strength in *X* while keeping the translator side comparatively stable. In the paired-data sweep, we varied the amount of data available to learn *h* with matched regularization. Representative learning curves are shown in Figure 8.

**Figure 8:**
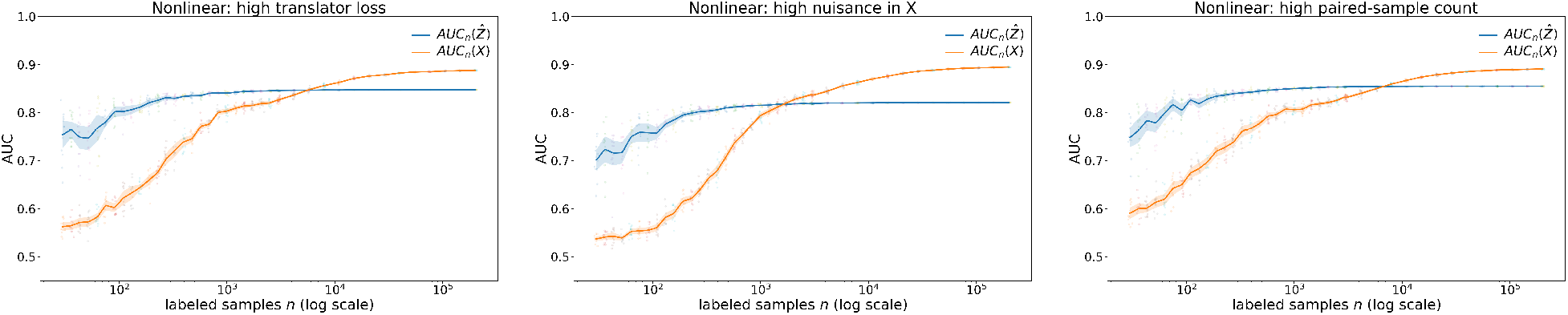
Representative nonlinear sweep families showing robust low-label advantage. Representative learning curves from three broader sweep families. **Left:** translator lossiness. **Middle:** nuisance entanglement in *X*. **Right:** paired-sample availability for learning *h*. In each case, 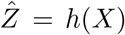 yields better performance in the low-label regime, whereas direct prediction from *X* improves more slowly and catches up only at larger labeled sample sizes.

**Figure 9:**
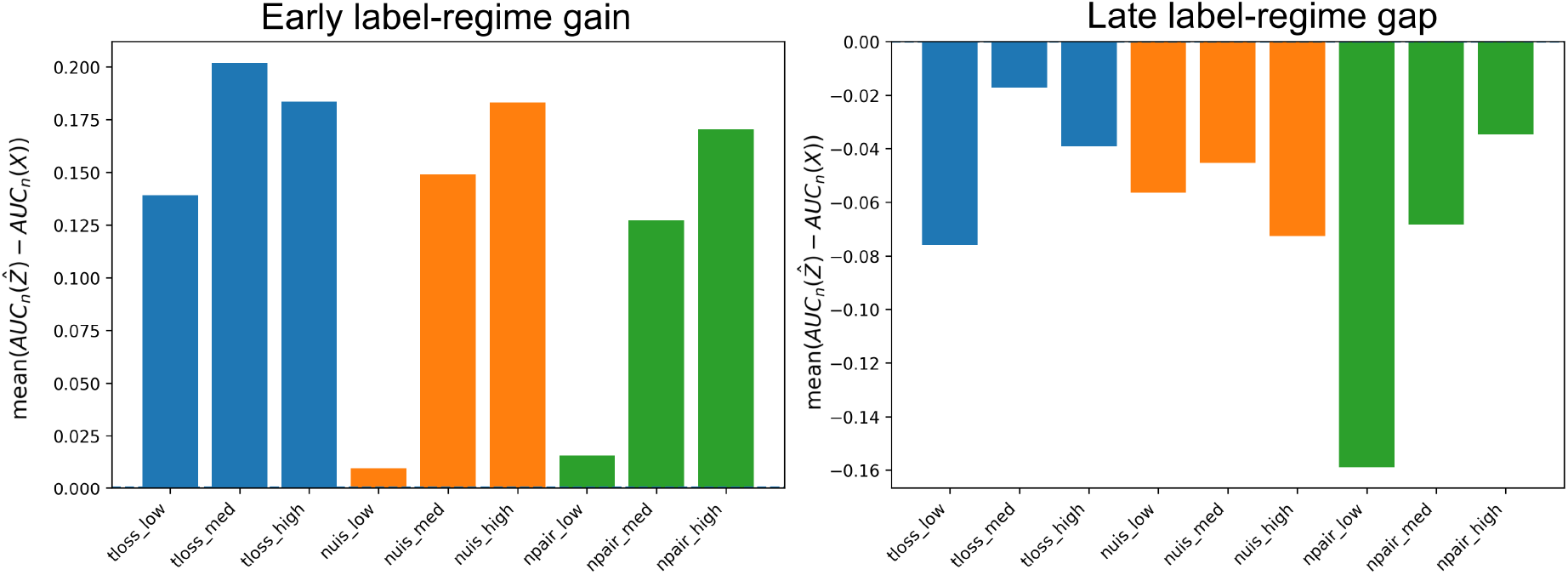
Finite-sample learnability advantage of translated representations in nonlinear regimes. Bars are grouped by experimental axis: translator lossiness (blue), nuisance strength in *X* (orange), and paired-sample count (green). **Left:** Δ_low-*n*_: all nonlinear conditions show positive gain, indicating improved learning efficiency under label scarcity. **Right:** Δ_tail_: values are negative across conditions, confirming that classifiers trained directly on *X* eventually match or slightly exceed 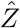 at larger label counts. Together the two panels show that translation improves finite-sample learnability without increasing asymptotic performance.

**Figure 10:**
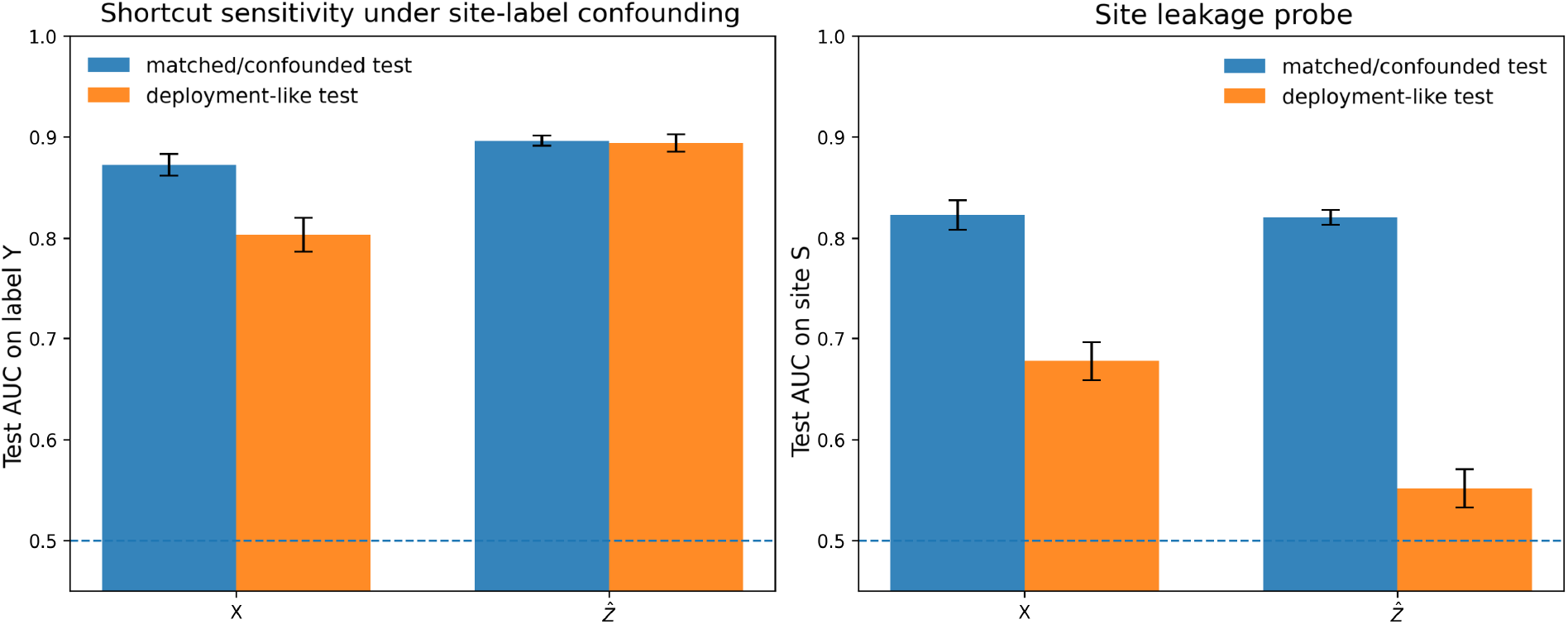
Site–label shortcut stress test with confounding removed at deployment-like evaluation. A translator 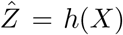 is learned from paired (*X, Z*^⋆^) data and then frozen. The paired cohort used to learn *h* is label-balanced but site-imbalanced, whereas downstream classifiers for *Y* are trained on labeled data with strong site–label confounding. The matched/confounded test preserves that correlation; the deployment-like test keeps site prevalence fixed but removes the site–label association. **Left:** test AUROC for predicting *Y* from *X* or 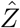. Direct prediction from *X* benefits from the matched/confounded setting and drops when the shortcut is broken, whereas prediction from 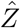 remains much more stable. **Right:** site-prediction probe, with chance shown at 0.5. Site is strongly recoverable from both representations under the matched/confounded mixture, but under the deployment-like mixture site predictability drops sharply, especially for 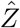, which approaches chance, indicating substantial suppression, though not guaranteed elimination, of shortcut structure.

To summarize across conditions more compactly than raw learning curves allow, we define the low-label gain and late-regime tail gap as

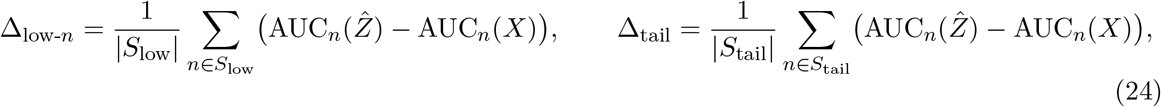

where *S*_low_ denotes the first six label-budget grid points and *S*_tail_ the final four.

Across all three sweep families, Δ_low-*n*_ *>* 0 and Δ_tail_ ≤ 0: the translated representation is systematically more effective in the low-label regime, while direct prediction from *X* eventually catches up or marginally surpasses it at the largest simulated label counts. The low-label advantage increases when the deployable representation is more strongly contaminated by nuisance and decreases when the translator is lossier or trained on fewer paired samples.

Across the linear benchmark, the nonlinear example, and the three sweep families, the conclusion is the same. Translation does not enlarge the deployment-time information set. What it can do is reorganize existing information into a representation with materially better finite-sample conditioning. The gain is strongest when the raw deployable representation is nuisance-entangled and labels are scarce, persists under nonlinear target structure, and can coexist with lower large-*n* performance when translation is sufficiently lossy.

### 3.5 False gains: confounding, miscalibration, and circularity

The preceding synthetic experiments isolate conditions under which deterministic translation can produce genuine finite-sample learnability gains under conserved deployment ceilings. That mechanistic result does not imply that an observed empirical gain in a real cohort should be interpreted as biology-aligned or deployment-relevant signal. In observational data, an apparent improvement in 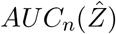 over *AUC*_*n*_(*X*) can also arise because the translated representation encodes shortcut structure correlated with the label, because it produces biologically plausible but poorly calibrated outputs, or because evaluation leaks information from the downstream test set into translator or hyperparameter selection. Accordingly, an observed gain 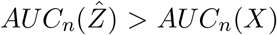 is necessary but not sufficient evidence that translation has closed a biologically meaningful method gap.

Three failure modes are especially important. First, **translator confounding**: 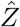 may retain or amplify site-, stain-, scanner-, or processing-specific structure that is correlated with *Y* in the labeled cohort. Second, **prior hallucination / poor calibration**: 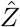 may appear biologically structured while remaining weakly calibrated to the underlying measured modality, for example by supporting confident downstream predictions in held-out settings where true signal should be absent or substantially attenuated. Third, **evaluation circularity**: apparent gains can be inflated when translator choice, preprocessing, corruption level, or downstream hyperparameters are selected using downstream test performance rather than a locked validation protocol. Site-stratified splits, frozen translator selection, controlled fidelity-degradation tests, and calibration against measured modalities when available are therefore required safeguards before treating gains from 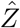 as biology-aligned.

This threat model also clarifies why the core proof-of-concept in Section 3 is synthetic. Real H&E cohorts do not permit the morphology-limited ceiling, structured nuisance, and translator fidelity to be varied independently while the underlying task is held fixed, so observed gains in such cohorts admit multiple explanations. The role of deployment-like evaluation is therefore not to negate the synthetic result, but to distinguish the learnability-mediated regime isolated above from shortcut-driven improvement in practice.

A minimal stress test introduces confounded nuisance and distribution mismatch. Let *S* denote site, allow nuisance components in *X* to correlate with *S*, and allow *S* to correlate with *Y* in the labeled cohort while the paired-data distribution used to learn *h*(·) is drawn from a different site mixture. In this setting, translation can (i) appear to improve random-split AUC by amplifying site-correlated structure, yet (ii) lose that advantage, or even reverse, under strict site-stratified or deployment-like evaluation. This pattern is diagnostic: it is more consistent with shortcut learning than with translation recovering a more biology-aligned representation, and it is closely aligned with the site-signature failure mode documented by (32).

## 4 TRACE and the Advantage Representation Curve (ARC)

Here we introduce TRACE (Translator Representation Analysis of Ceilings and Efficiency), an open-source toolkit (https://github.com/psaisan/TRACE) that operationalizes the formal framework developed above. TRACE is designed to characterize and diagnose biomarker prediction performance from direct learning on *X* versus translated learning on *h*(*X*), to analyze paired learning curves, and to generate theory-driven simulations. Beyond comparing direct deployable-representation prediction against a single translated representation, TRACE is meant to help evaluate and navigate a growing family of deployed translators in terms of learnability, robustness, and likely ceiling-limited failure.

Central to TRACE’s analytics is a diagnostic function defined as follows. Let

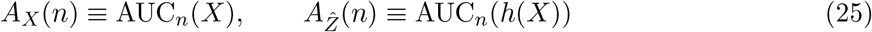

denote the expected test AUROC of a fixed downstream learner trained on *n* labeled samples using, respectively, the deployable representation *X* and the translated representation *h*(*X*). Both curves are defined relative to a fixed downstream protocol; ARC is therefore a property of the representation-plus-protocol pair, not of the representation alone. The *Advantage Representation Curve* (ARC) is then

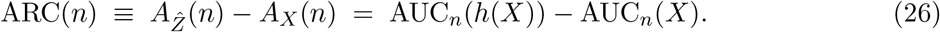

ARC(*n*) encodes the expected generalization performance gap, as a function of label scale, between models trained on *h*(*X*) and on *X*. Its role is to reveal where performance gains are concentrated, whether and when direct learning from *X* catches up, and whether the translated representation becomes negligible or harmful.

ARC(*n*) also admits a precise interpretive decomposition in terms of deployment ceilings and method gaps. Define the *method gap* for a representation *R* at label budget *n* as

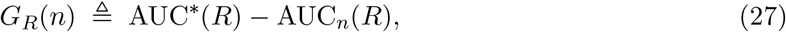

where AUC^*^(*R*) denotes the Bayes-optimal deployment ceiling for *R*. Note that *G*_*R*_(*n*) combines finite-*n* estimation effects with any persistent mismatch between the downstream learner class and the Bayes-optimal classifier; it is not a purely finite-sample quantity unless the downstream learner is Bayes-consistent in *R*. Then

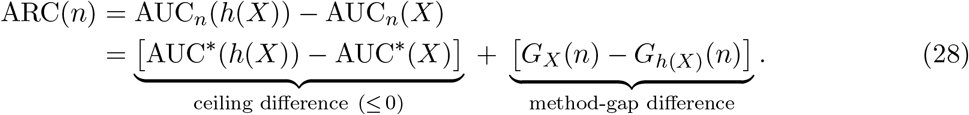

The ceiling difference is non-positive under deterministic deployed translation, since AUC^*^(*h*(*X*)) ≤ AUC^*^(*X*). The method-gap difference is positive when the translated representation is easier to learn from at label budget *n* under the chosen protocol. Positive ARC therefore does not mean that translation raises the deployment ceiling; it means that method-gap reduction outweighs any ceiling loss incurred by translation. Equation (28) is an interpretive decomposition rather than a directly observed finite-sample quantity: TRACE estimates ARC(*n*) from resampled learning curves and uses the ceiling/method-gap split as a lens for interpreting curve shape, not as two separately measurable components.

### 4.1 Canonical ARC regimes and summary statistics

TRACE organizes observed ARC curves into four canonical regimes that provide its main diagnostic vocabulary:

- **Quick-gain:** ARC is positive at low *n* and decays toward zero as direct learning from *X* catches up.
- **Sustained-gain:** ARC remains positive across the full studied label range.
- **Neutral:** ARC remains near zero; neither representation has a consistent advantage.
- **Impaired:** ARC is negative or reverses sign; translation underperforms direct learning from *X*.

The boundaries between regimes are not rigid; real studies can lie between them, and mixed cases are reflected as such rather than forced into artificially pure regime calls.

To move from the full ARC curve to a compact regime score, TRACE computes three summary statistics. The first two capture early and late label-regime behavior; the third locates the scale at which any initial advantage disappears:

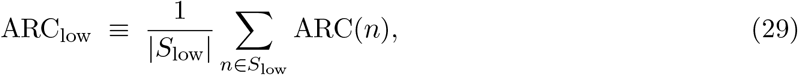

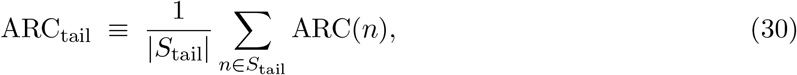

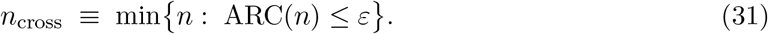

By default, TRACE defines *S*_low_ and *S*_tail_ as the first and last quartiles of the user-specified label grid, and reports *n*_cross_ as the first crossing of a smoothed ARC(*n*) below *ε* = 0; cases in which ARC never crosses *ε* on the studied grid are right-censored at *n*_max_. The regime-score panel is then computed by soft thresholding on (ARC_low_, ARC_tail_, *n*_cross_) against reference templates for each canonical regime, returning a similarity profile rather than a hard classification. These summaries do not replace the full curve; they provide the quantitative basis for the regime-score panel shown in Figure 11.

**Figure 11:**
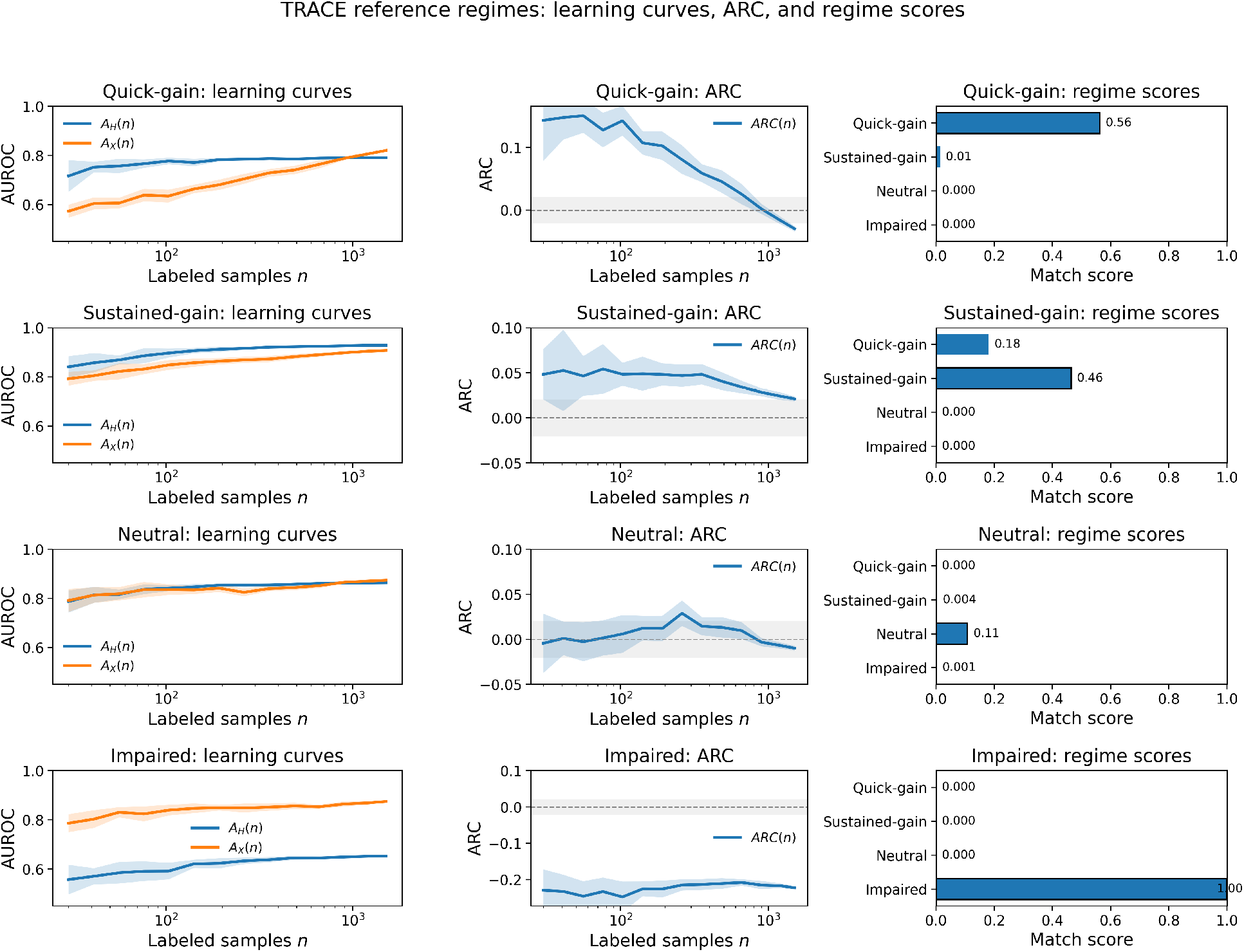
TRACE reference outputs across four canonical ARC regimes. **Left:** Each row shows paired learning curves 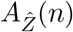 and *A*_*X*_(*n*). **Middle:** the corresponding advantage representation curve 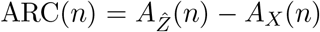. **Right:** regime-score similarity panel. **Top to bottom:** quick-gain, sustained-gain, neutral, and impaired. The regime scores quantify similarity to reference templates rather than probabilities, allowing mixed or borderline cases to be represented without forcing a hard class label.

For a given study, TRACE returns three coordinated outputs: (i) paired learning curves *A*_*X*_(*n*) and 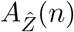 with uncertainty bands, (ii) the ARC curve with uncertainty, and (iii) a regime-score panel quantifying similarity to the canonical ARC regimes.

In this paper, the theory identifies the relevant signatures; the controlled synthetic experiments isolate their mechanism; TRACE provides the reusable workflow that measures those signatures in a study-ready form. TRACE is the diagnostic framework, and ARC is its central empirical readout of translator-induced finite-sample advantage under conserved deployment ceilings.

## 5 Clinical deployment at a translational inflection point

Genomic profiling remains the standard for targeted treatment selection in oncology (17; 16). In practice, however, comprehensive biomarker testing in NSCLC can be constrained by limited tissue, sample adequacy, and turnaround time, especially for small biopsies (19). These constraints motivate decision-support approaches that extract useful signal from routine H&E while preserving tissue for confirmatory assays and broader profiling, including liquid biopsy when appropriate (22; 18; 20; 21).

From a clinical perspective, the dominant question is often not whether morphology contains any signal, but whether that signal can be learned robustly under realistic label budgets and heterogeneity. Empirically, weakly supervised whole-slide learning reaches near-clinical performance for morphology-dominant tasks (1), whereas mutation prediction from H&E remains more variable and more sensitive to cohort shift (2; 13; 14; 15; 25; 26). This is exactly the regime in which our distinction between modality-limited ceilings and method-limited gaps becomes clinically relevant.

Deployed translation systems such as virtual spatial transcriptomics or virtual protein mapping do not add new slide-specific measurements at inference and therefore cannot raise the per-slide H&E information ceiling. Their practical value is instead that they can encode population-level morphology-to-biology structure learned from paired data, potentially improving robustness under realistic heterogeneity by suppressing nuisance and exposing biology-aligned axes (4; 3). In the near term, the most realistic role of such systems is triage and decision support rather than replacement of confirmatory molecular assays. Clinically, the relevant readout is therefore not only mean AUC, but reliability under deployment-like evaluation. If translation is truly closing a method-limited gap, improvements should concentrate in low-label regimes, increase under heterogeneity, and degrade predictably as translation fidelity decreases. Gains that vanish under site-stratified evaluation or fail fidelity-degradation tests are more consistent with shortcut learning than with clinically meaningful, biology-aligned improvement (26; 15).

## 6 Discussion

The empirical trajectory of histology-based molecular prediction has largely been a race of scale; larger cohorts, deeper encoders, stronger multiple-instance learning, and increasingly elaborate heuristics. Yet for many targets, the field repeatedly reconverges on similar performance plateaus. High-fidelity molecular translators such as virtual spatial transcriptomics and protein mapping offer a powerful new avenue to engage those plateaus, but they also sharpen a distinction that is easy to blur in practice: a measured molecular assay introduces new slide-specific information at inference, whereas a deployed deterministic translator does not.

We make that distinction explicit by decoupling the H&E deployment ceiling from the finite-sample method gap of weakly supervised learning. Translators do not raise the ceiling. Rather, they import population-level morphology–biology structure learned from paired cohorts and reorganize existing H&E signal into a representation that can be easier to learn from under label scarcity, structured nuisance, and cohort heterogeneity. This explains how 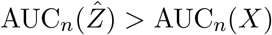 can arise in practice even when 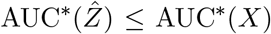: the gain is not new slide-specific information, but leaps in learnability inherited from rich paired molecular supervision that trained the translator.

This reframing changes what counts as evidence. Gains that concentrate under limited labeled data, strengthen under heterogeneity, and deteriorate as translation fidelity degrades are consistent with a removable method gap. By contrast, when both direct and translated performance flatten, and additional labels or improvements in translation no longer meaningfully change the curves, after shortcut-driven and evaluation-artifact explanations have been excluded, the remaining constraint is more consistent with the measurement modality itself. TRACE and ARC formalize this logic as a practical way to decide whether to invest next in labels, in paired supervision, in robustness, or in a richer representation.

The broader implication is that translators play a dual role in computational pathology. In method-limited regimes they are interventions: they make existing morphology more learnable. As that gap narrows, they become unusually demanding probes of what H&E can and cannot support, precisely because they are trained against rich paired molecular supervision rather than sparse endpoint labels alone. When strong translators still fail to unlock further gains under deployment-aware evaluation, the result is not merely another negative benchmark. It is evidence that the remaining limitation may lie less in the learner than in the modality.

That is also why the framework points forward rather than inward. The ceiling–gap vocabulary developed here is not specific to H&E or to any single translator; it offers a decision logic for the next generation of biomarker pipelines. In some settings, H&E will remain the primary substrate, with translators improving triage, enrichment, and robustness. In others, the more productive path will be deliberate fusion with more informative yet increasingly practical modalities—targeted stains, multiplexed imaging, spatial transcriptomics, proteomic mapping, or selective hybrid workflows that reserve costly assays for cases in which morphology has become information-limited. None of this diminishes what H&E-based computational pathology has achieved. Extracting molecular signal from a century-old staining protocol is a remarkable feat, and H&E prediction will retain an essential clinical role for the foreseeable future. What changes is that the ceiling is no longer something to lament or overlook. It is now something to measure, and measuring it is what transforms a recurrent plateau from a mystery into a decision.

## Footnotes

1 Data processing inequality (DPI) for deterministic functional transforms (27). AUROC: finite-*n* (fixed pipeline); no MI–AUROC monotonicity assumed.

2 Learnability (finite-*n*): under a fixed protocol and labeled training set size *n, R*_1_ ≻ *R*_2_ iff AUC_*n*_(*R*_1_) *>* AUC_*n*_(*R*_2_). This notion is distinct from Bayes-optimal deployment ceilings.

## References

[1] G. Campanella, M. G. Hanna, L. Geneslaw, et al. Clinical-grade computational pathology using weakly supervised deep learning on whole slide images. Nature Medicine, 25(8):1301–1309, 2019. doi:10.1038/s41591-019-0508-1.

[2] N. Coudray, P. S. Ocampo, T. Sakellaropoulos, et al. Classification and mutation prediction from non-small cell lung cancer histopathology images using deep learning. Nature Medicine, 24(10):1559–1567, 2018. doi:10.1038/s41591-018-0177-5.

[3] J. M. J. Valanarasu, H. Xu, N. Usuyama, et al. Multimodal AI generates virtual population for tumor microenvironment modeling. Cell, 189(2):386–400.e19, 2026. doi:10.1016/j.cell.2025.11.016.

[4] B. Schmauch, L. Herpin, A. Olivier, et al. A deep learning-based multiscale integration of spatial omics with tumor morphology. Nature Communications, 16(1):11674, 2025. doi:10.1038/s41467-025-66691-y.

[5] Y. Şenbabaoğlu, V. Prabhakar, A. Khormali, et al. MOSBY enables multi-omic inference and spatial biomarker discovery from whole slide images. Scientific Reports, 14(1):18271, 2024. doi:10.1038/s41598-024-69198-6.

[6] D.-T. Hoang, G. Dinstag, E. D. Shulman, et al. A deep-learning framework to predict cancer treatment response from histopathology images through imputed transcriptomics. Nature Cancer, 5(9):1305–1317, 2024. doi:10.1038/s43018-024-00793-2.

[7] V. Vapnik and A. Vashist. A new learning paradigm: Learning using privileged information. Neural Networks, 22(5–6):544–557, 2009. doi:10.1016/j.neunet.2009.06.042.

[8] T. G. Dietterich, R. H. Lathrop, and T. Lozano-Pérez. Solving the multiple instance problem with axis-parallel rectangles. Artificial Intelligence, 89(1–2):31–71, 1997. doi:10.1016/S0004-3702(96)00034-3.

[9] M. Ilse, J. M. Tomczak, and M. Welling. Attention-based deep multiple instance learning. In Proceedings of the 35th International Conference on Machine Learning, volume 80 of Proceedings of Machine Learning Research, pages 2127–2136, 2018.

[10] M. Gadermayr and M. Tschuchnig. Multiple instance learning for digital pathology: A review of the state-of-the-art, limitations & future potential. Computerized Medical Imaging and Graphics, 112:102337, 2024. doi:10.1016/j.compmedimag.2024.102337.

[11] D. Lopez-Paz, L. Bottou, B. Schölkopf, and V. Vapnik. Unifying distillation and privileged information. In International Conference on Learning Representations (ICLR), 2016.

[12] J. N. Kather, A. T. Pearson, N. Halama, et al. Deep learning can predict microsatellite instability directly from histology in gastrointestinal cancer. Nature Medicine, 25(7):1054–1056, 2019. doi:10.1038/s41591-019-0462-y.

[13] W. Zhang, W. Wang, Y. Xu, et al. Prediction of epidermal growth factor receptor mutation subtypes in non-small cell lung cancer from hematoxylin and eosin-stained slides using deep learning. Laboratory Investigation, 104(8):102094, 2024. doi:10.1016/j.labinv.2024.102094.

[14] J. J. Pao, M. Biggs, D. Duncan, et al. Predicting EGFR mutational status from pathology images using a real-world dataset. Scientific Reports, 13(1):4404, 2023. doi:10.1038/s41598-023-31284-6.

[15] M. H. Nguyen, M. H. N. Le, A. T. Bui, and N. Q. K. Le. Artificial intelligence in predicting EGFR mutations from whole slide images in lung cancer: a systematic review and meta-analysis. Lung Cancer, 204:108577, 2025. doi:10.1016/j.lungcan.2025.108577.

[16] J.-C. Soria, Y. Ohe, J. Vansteenkiste, et al. Osimertinib in untreated EGFR-mutated advanced non-small-cell lung cancer. New England Journal of Medicine, 378(2):113–125, 2018. doi:10.1056/NEJMoa1713137.

[17] G. J. Riely, D. E. Wood, D. L. Aisner, et al. NCCN Guidelines® Insights: Non-Small Cell Lung Cancer, Version 7.2025. Journal of the National Comprehensive Cancer Network, 23(9):354–362, 2025. doi:10.6004/jnccn.2025.0043.

[18] D. Di Capua, D. Bracken-Clarke, K. Ronan, A.-M. Baird, and S. Finn. The liquid biopsy for lung cancer: state of the art, limitations and future developments. Cancers, 13(16):3923, 2021. doi:10.3390/cancers13163923.

[19] U. Chaddha, A. Agrawal, U. Ghori, et al. Safety and sample adequacy for comprehensive biomarker testing of bronchoscopic biopsies: an American Association of Bronchology and Interventional Pulmonology and International Association for the Study of Lung Cancer clinical practice guideline. Journal of Thoracic Oncology, 20(9):1237–1256, 2025. doi:10.1016/j.jtho.2025.05.014.

[20] D. Y. Zhang, A. Venkat, H. Khasawneh, et al. Implementation of digital pathology and artificial intelligence in routine pathology practice. Laboratory Investigation, 104(9):102111, 2024. doi:10.1016/j.labinv.2024.102111.

[21] M. G. Hanna and O. Ardon. Digital pathology systems enabling quality patient care. Genes, Chromosomes & Cancer, 62(11):685–697, 2023. doi:10.1002/gcc.23192.

[22] L. E. Raez, K. Brice, K. Dumais, et al. Liquid biopsy versus tissue biopsy to determine front line therapy in metastatic non-small cell lung cancer (NSCLC). Clinical Lung Cancer, 24(2):120–129, 2023. doi:10.1016/j.cllc.2022.11.007.

[23] G. Campanella, N. Kumar, S. Nanda, et al. Real-world deployment of a fine-tuned pathology foundation model for lung cancer biomarker detection. Nature Medicine, 31(9):3002–3010, 2025. doi:10.1038/s41591-025-03780-x.

[24] J. Park, S. Shin, W. Hwang, et al. Deep learning predicts EGFR mutation status from histology images in non-small cell lung cancer. Cancer Research Communications, 5(12):2127–2141, 2025. doi:10.1158/2767-9764.CRC-25-0155.

[25] Y. Zhao, S. Xiong, Q. Ren, et al. Deep learning using histological images for gene mutation prediction in lung cancer: a multicentre retrospective study. The Lancet Oncology, 26(1):136–146, 2025. doi:10.1016/S1470-2045(24)00599-0.

[26] C. Rolfo, E. Ofek, I. Barshack, et al. Validation of histopathology-based deep learning algorithms for detection of actionable non-small cell lung cancer biomarkers. npj Precision Oncology, 10(1):62, 2026. doi:10.1038/s41698-025-01267-z.

[27] T. M. Cover and J. A. Thomas. Elements of Information Theory. Wiley-Interscience, 2nd edition, 2006.

[28] D. Ruiz, P. Cárdenas, L. Manrique, D. Vega, G. M. Mejia, and P. Arbeláez. Completing spatial transcriptomics data for gene expression prediction benchmarking. Medical Image Analysis, 106:103754, 2025. doi:10.1016/j.media.2025.103754.

[29] The Cancer Genome Atlas Research Network. Comprehensive molecular profiling of lung adenocarcinoma. Nature, 511(7511):543–550, 2014. doi:10.1038/nature13385.

[30] Y. H. Wen, E. Brogi, A. Hasanovic, M. Ladanyi, R. A. Soslow, D. Chitale, J. Shia, and A. L. Moreira. Immunohistochemical staining with EGFR mutation-specific antibodies: high specificity as a diagnostic marker for lung adenocarcinoma. Modern Pathology, 26(9):1197–1203, 2013. doi:10.1038/modpathol.2013.53.

[31] A. Warth, R. Penzel, H. Lindenmaier, et al. EGFR, KRAS, BRAF and ALK gene alterations in lung adenocarcinomas: patient outcome, interplay with morphology and immunophenotype. European Respiratory Journal, 43(3):872–883, 2014. doi:10.1183/09031936.00018013.

[32] F. M. Howard, J. Dolezal, S. Kochanny, et al. The impact of site-specific digital histology signatures on deep learning model accuracy and bias. Nature Communications, 12(1):4423, 2021. doi:10.1038/s41467-021-24698-1.

[33] B. Schmauch, A. Romagnoni, E. Pronier, et al. A deep learning model to predict RNA-seq expression of tumours from whole slide images. Nature Communications, 11(1):3877, 2020. doi:10.1038/s41467-020-17678-4.

[34] Y. Zeng, Z. Wei, W. Yu, et al. Spatial transcriptomics prediction from histology jointly through Transformer and graph neural networks. Briefings in Bioinformatics, 23(5):bbac297, 2022. doi:10.1093/bib/bbac297.

[35] Y. Jia, J. Liu, L. Chen, T. Zhao, and Y. Wang. THItoGene: a deep learning method for predicting spatial transcriptomics from histological images. Briefings in Bioinformatics, 25(1):bbad464, 2024. doi:10.1093/bib/bbad464.

[36] L. Farndale, R. Insall, and K. Yuan. TriDeNT: Triple deep network training for privileged knowledge distillation in histopathology. Medical Image Analysis, 102:103479, 2025. doi:10.1016/j.media.2025.103479.

[37] G. Alain and Y. Bengio. Understanding intermediate layers using linear classifier probes. In International Conference on Learning Representations (ICLR) Workshop Track, 2017.

